# Plant origin determines seed mass, seed nutrients and germination behavior of a dominant grass species

**DOI:** 10.1101/2020.03.02.973552

**Authors:** Andrea Veselá, Lien Duongová, Zuzana Münzbergová

**Author notes:** Both authors contributed equally (Andrea Veselá, Lien Duongová).

## Abstract

Although number of studies exploring effects of climate change on plants is increasing, only few studies pay attention to germination. Understanding of germination behaviour is complicated by impact of climate on seed mass and possibly also seed nutrients, which play irreplaceable role in nourishing the sprout. The germination behaviour of species may also depend on trade-off between generative and vegetative reproduction.

We studied *Festuca rubra* originating from localities situated along a natural climatic grid. Seeds of different origin were germinated in two temperature and two moisture regimes, simulating the extremes of the grid. To study relationship between generative and vegetative reproduction we used data on vegetative traits originating from the same study system.

Seed mass and nutrient concentrations (nitrogen and phosphorus) were significantly affected by original climate, while carbohydrates not. Higher seed mass and warm temperature of origin caused higher and faster germination. Warm and dry condition during germination caused the lowest germination but the highest seed viability. Total germination and proportion of viable seeds negatively correlated with plant performance variables contributing to vegetative reproduction. Despite this, the patterns detected using datasets of germination and plant performance, did not provide fully comparable results.

Simulated changes in climatic condition can modify seed mass and seed nutrients and these changes translate into changes in species germination behavior. After accounting for seed mass, both original and target conditions determine species germination indicating that both genetic differentiation as well as actual conditions drive the germination patterns. As the patterns detected at the level of seed germination do not fully match those detected for the vegetative traits, we urge that future studies should focus on multiple stages of plant life to understand species responses to future climates.

## Introduction

One of the most commonly observed responses to climate change is migration, often to higher altitudes (Jump and Penuelas 2005). However, species are not always able to migrate (Jump and Penuelas 2005) due to slow migration rates (Huntley 1991) (Neilson et al. 2005) or lack of habitats suitable for migration (e.g. in alpine species already at the peak of mountains). Species thus need to use other strategies to respond to the changes.

One possibility is the response through phenotypic plasticity, which involves expression of different morphology or physiology by plants of the same genotype under different environment. It can be observed e.g. in seed traits (including germination), plant height or flowering time (Nicotra et al. 2010) (Anderson et al. 2012) (Wainwright and Cleland 2013) (Anderson and Gezon 2015). This mechanism allows species to cope mainly with short-term undirected changes. Alternatively, species may respond by genetic adaptation resulting from climate selecting individuals with traits advantageous for survival in the specific conditions. Although genetic adaptation is usually slower, it allows the species to respond to stronger directional changes occurring over longer time periods. Despite the increasing number of studies assessing the importance of these two mechanisms to changing climatic conditions (e.g. (Jump and Penuelas 2005) (Wilczek et al. 2010)), our knowledge on their importance in species germination behaviour is still rather limited (Nicotra et al. 2010).

Paying special attention to germination is important as early developmental stages of plants are expected to be more sensitive to climate change than adult stages (Lloret, Penuelas and Estiarte 2004) (Fay and Schultz 2009) (Dalgleish, Koons and Adler 2010). Germination is mainly affected by light, temperature and water availability (Baskin and Baskin 2001, Fenner and Thompson 2005) and may thus be strongly affected by ongoing climate change. The response of seed germination to temperature has been extensively studied (e.g.(Grime et al. 1981), (Schütz and Rave 1999) (Gardarin, Daurr and Colbach 2011)), studies exploring germination response to variation in moisture are less common (but see e.g. (Wen et al. 2015) (Ruhl et al. 2015). In addition, some studies demonstrated interaction of temperature and moisture on plant performance (e.g. (Suseela et al. 2012) (Meineri, Spindelbock and Vandvik 2013) (Münzbergová et al. 2017). There are studies of these interactive effects on germination using species from arid regions (e.g. Rivas-Arancibia et al. 2006, Gurvich et al. 2017, (Flores, Perez-Sanchez and Jurado 2017)) and species of alpine conditions, likely being most threatened by the ongoing change in climate, are usually neglected (but see (Orsenigo et al. 2015), Veselá et al. submitted). Both studies indicate decreasing germination in warm, dry conditions.

Germination response may depend not only on current conditions, but can also differ intra-specifically due to species adaptation to the climate of origin (Meyer, Allen and Beckstead 1997) (Qaderi and Cavers 2002) (Degreef et al. 2002). Many studies, for example, demonstrated that populations of a single species differ in germination percentage with populations coming from the warmest conditions having the highest germination (Cruz et al. 2003) (Ndihokubwayo, Nguyen and Cheng 2016) (Santo, Mattana and Baechetta 2015) (Bauk et al. 2017) (Mira, Arnal and Perez-Garcia 2017). In addition, the differences between seeds of different origin may depend on the actual conditions in which the seeds are produced (Veselá at al. submitted).

Effect of seed origin on germination response could be also caused by different seed mass and nutrient content in the seeds because favorable conditions can contribute to higher seed reserve and thus to higher seed mass. Relationship between seed mass and germination is commonly studied. Positive effect of seed mass on germination was demonstrate in studies of (Münzbergová and Plačková 2010) (Wu and Du 2007) and (Paulů, Harčariková and Münzbergová 2017), but see e.g. (Wang et al. 2009) (Wu, Li and Du 2011) for an opposite trend. Studies focus seed nutrients are less frequent, but (Bu et al. 2018), for instance, demonstrated differences between populations in the content of nitrogen and phosphorus, with higher content of both nutrients in alpine conditions. Carbohydrates, nitrogen and phosphorus provide the initial energy and nutrients for growth of the sprout (Muthukumar and Udaiyan 2000) (McGinley and Charnov 1988) and the differences in their content may thus drive the differences in seed germination behavior. (Rees et al. 2001) suggested that heavier seeds have higher content of nutrients, which should contribute to their higher germination, but did not quantitatively demonstrate the relationship between nutrients and seed mass. In addition, intra-specific variability of nutrient concentration in seeds is not sufficiently explored (but see (Vaughton and Ramsey 2001) (Obeso 2012) (Kolodziejek 2017)) and to our best knowledge, the effects of nutrient concentration on germination behavior have not been studied yet.

Despite the increasing bulk of knowledge on the determinants of species germination (Bauk et al. 2017) (Veselá et al. submitted) and species performance (Meineri et al. 2013) (Münzbergová et al. 2017) in response to climate, our knowledge on the correspondence of the patterns of responses between the seeds and grown up plants in long-lived species under climate change is largely missing. Understanding the effects of changes in climatic conditions on multiple parts of life cycle is, however, important as the patterns detected in single life cycle stages may contradict the conclusions based on other stages (Münzbergová 2005) (Kolb, Dahlgren and Ehrlen 2010) (Laughlin et al. 2018). In natural conditions, several studies have detected negative correlation between generative and vegetative reproduction (e.g. (Cheplick 1995) (Worley and Harder 1996) (Ronsheim and Bever 2000) (van Kleunen, Fischer and Schmid 2002) (Herben et al. 2012)), which indicate species adaptation on environment and preference more advantageous reproduction. However, other studies did not find similar correlation (e.g. (Reekie 1991) (Cain and Damman 1997)). This may suggest that patterns detected at the stage of generative reproduction may go in the opposite direction than patterns detected for vegetative reproduction. To what extend the mechanisms driving these two types of responses to climate change correspond to each other and to what extend the values of the different traits correlate, however, remains largely unexplored.

The aim of this study is to understand the importance of original and actual climatic conditions, and thus the importance of genetic differentiation and phenotypic plasticity, for species germination using natural factorially crossed temperature and moisture gradients and widespread clonal grass species, *Festuca rubra*, as the model system. Each locality represent different type of climate and other factors (e.g. slope, bedrock, grazing history) are similar as much as possible (Meineri et al. 2014). Seed mass and nutrient content in the seeds may be an important mechanism affecting species germination patterns and at the same time, they may be affected by species origin. We thus studied the effect of plant origin on seed mass and nutrient content and the effect of seed mass and nutrient content on seed germination in different climatic conditions further referred as a target condition. To assess to what extend the effects detected at the level of germination correspond to the effects detected using grown-up plants, we used vegetative data from the same study system published in (Münzbergová et al. 2017) and compared the patterns to the results obtained here.

Specifically, we aimed to answer the following questions: i) Does the original climate affect seed nutrient content and seed mass and do these differences, if any, relate to seed germination patterns?, ii) What is the effect of original and target conditions on species germination patterns and do these factors interact?, iii) What is the relationship between traits related to generative and vegetative reproduction and their responses to changing climatic conditions?

We tested the following hypotheses: i) Seed mass and seed nutrient content, especially the content of nitrogen and phosphorus, will be affected by original climate, with the highest nitrogen and phosphorus content in the alpine populations. Higher seed mass will be caused mainly by higher content of carbohydrates, because these compounds have higher molecular weight than nitrogen and phosphorus. Simultaneously, higher seed mass and higher concentration of nutrients, especially carbohydrates, will contribute to higher germination., ii) Target temperature and moisture will interact in their effect on seed germination, and germination percentage will decrease with increasing temperature and decreasing moisture. We also expected that germination response will be strongly affected by origin of the population and the highest germination percentage will be observed in populations coming from the warmest conditions. Also, origin and target conditions will interact. iii) Total germination and/or proportion of viable seeds will negatively correlate with plant performance variables contributing to vegetative reproduction. Factors determining species germination patterns will be the same as the factors determining vegetative reproduction, just acting in the opposite direction.

## Materials and methods

### Study species and localities

We chose *Festuca rubra* L., a widespread perennial grass species of temperate grasslands in Europe, as a model plant. In the experiment, we used a widespread hexaploid type from the *F. rubra* complex (Šurinová et al. 2019). It reproduces by seeds as well as vegetatively, producing both intravaginal and extravaginal tillers on rhizomes. It grows at different densities in grasslands, both as a dominant with only a few other species and also as a subordinate of species-rich stands. *F. rubra* possesses considerable genetic variability and plasticity and can adapt to a wide range of climatic conditions (Skalova et al. 1997) (Herben et al. 2001) (Münzbergová et al. 2017) (Münzbergová and Hadincová 2017).

The experimental plants originated from localities occurring in a unique natural climatic grid in western Norway previously used e.g. by (Meineri et al. 2014) and (Klanderud, Vandvik and Goldberg 2015) as we wanted to have plants from defined environmental conditions. The grid is represented by twelve grassland localities (Fig. 1) combining four levels of mean annual precipitation [ca. 600 (1), 1300 (2), 2000 (3) and 2700 (4) mm/year] and three levels of mean summer temperature [defined as the mean of the four warmest months; ca 6.5°C (alpine, ALP), 8.5°C (sub-alpine, SUB) and 10.5°C (boreal, BOR). *F. rubra* occurs on 11 localities in climatic grid and does not occur on the twelfth locality (ALP2). Localities were selected specifically to ensure that grazing regime and grazing history, bedrock, slope, aspect and vegetation types are as similar as possible (Meineri et al. 2014). The communities are grazed intermediate-rich meadows (Potentillo-Festucetum ovinae; G8 *sensu* Fremstad 1997) occurring on south-west facing (with the exception LOW3, which was exposed to the east), shallow slope (5-20°) with relatively rich bedrock in terms of nutrient availability. Geographical distance between the sites ranges from 0.65 km to 175 km (BOR1 and BOR4). The two geographically closest localities, BOR2 and SUB2, which are only 0.65 km apart are 400 m in altitude, and hence differ substantially in climate (Meineri et al. 2014).

**Fig 1.**
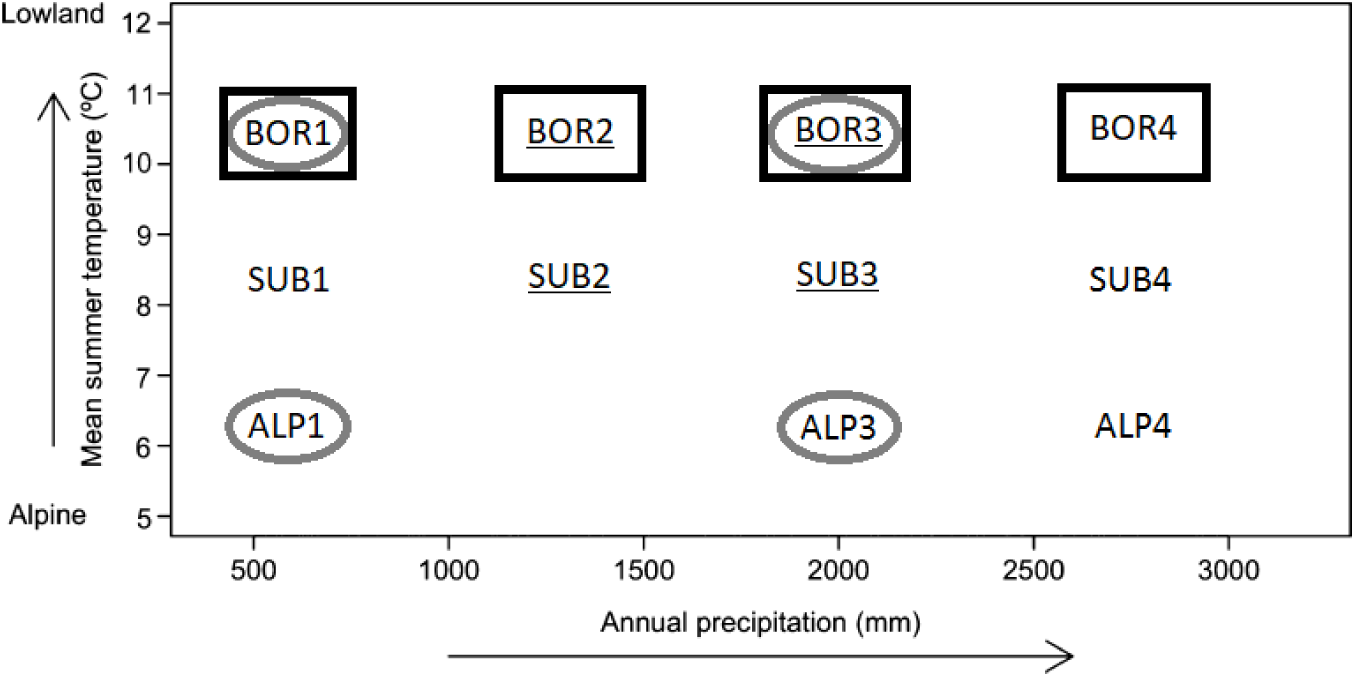
Position of the localities in the climate grid. All shown populations were included into the first analysis. The populations included into the second analysis are highlighted by square, the populations included into the third analysis are underlined and populations included in nutrient analysis are circled.

### Plant material and seed collection

For this study, we used plants collected for the purpose of our previous studies (Münzbergová and Hadincová 2017, Münzbergová et al. 2017) (Knappová et al. 2018) (Münzbergová et al. 2018). Specifically, the material consisted of 40 *F. rubra* plants collected in July 2014, being at least 1 m apart from each other at each locality. Living plants were transported to the experimental garden of the Institute of Botany, Czech Academy of Sciences in Průhonice, Czech Republic (49°59’38.972”N, 14°33’57.637”E; mean temperature of the four warmest months 16.5°C and regular watering during the vegetation season) and planted into pots (16 × 16 × 16 cm, filled with a mixture of common garden soil and sand in 1:2 ratio) just after the transport at the end of July 2014. The common garden soil comprised compost from the experimental garden containing approximately 0.135 % of nitrogen, 1.35% of carbon and 46.5 mg of phosphorus in 1000 g of soil. After the plants recovered from the transport, they were extracted from the pots and reduced to a single ramet. This was done to make sure that we only have one genotype per pot avoiding the possibility that we originally collected multiple intermingled genotypes.

Plants from all 12 populations were cultivated for 2 years before collecting the seeds. In early spring 2016, the plants, that were about to flower were transferred into 120 × 80 × 60 cm cages covered with a double layer of fine mesh fabric. The mesh size was enabling the light and wind to go through the cage, but preventing the pollen from outside to penetrate in. In each cage, only plants from one locality were placed. In this way, the plants could be cross-pollinated within population but not between populations. From July to August 2016 we collected the ripe seeds and kept the seeds of each mother plant separately. The seeds were stored at room temperature (about 20°C and ambient moisture of about 55%) for 10 weeks before the experiments. All plants performed well in the experimental garden, and it is thus unlikely that quality of the seeds would be affected by low performance of some of the plants in the garden. We thus expect that differences in seed germination behavior are not due to different distance between the natural sites and the experimental garden, but rather due to effect of the original conditions (see discussion for more details on the issue).

### Nutrient analyses

To assess whether content of nutrients depends on original climatic conditions, we determined the content of nitrogen, phosphorus and carbohydrates in the form of starch and fructans in seeds from 4 populations (circled in Fig 1.). While we initially planned to used seeds from the 4 most extreme populations, we had to replace seeds from ALP4 and BOR4 with seeds from ALP3 and BOR3 because of insufficiency of seeds from ALP4. To assess whether seed nutrients differ between mother plants, we determined the content of nutrients in 10 randomly chosen mother plant from each population (subset of plants used for the germination test) i.e. we analyzed 40 mothers from 4 populations. While we initially intended to analyze the nutrients in the remaining populations in the next step, we did not do this as this initial testing showed strong correlation of all the nutrients with seed mass (Supporting information 1).

Content of nitrogen was determined according the Kjeldal methodology (Kjeldahl 1883), content of phosphorus was determined by mineralization by perchloric acid (HClO_4_) (Talvitie, Illustre and Perez 1962). Content of starch and fructans was measured by enzymatic procedure Megazyme (McCleary et al. 1994). All analyses of all mothers included 3 replicates. The nutrient analyses were performed in the Analytic Laboratory of the Institute of Botany of the Czech Academy of Sciences. All nutrients were expressed as a percentage of dry biomass.

### Seed mass

For each population (11), we randomly selected at least twenty mother plants, for which we determined weight of all collected seeds of one mother plant and number of seeds in the sample. These values were used to express weight of thousand seeds.

### Germination tests

From each population (11), we chose 14 to 20 mother plants which produced at least 20 undamaged, fully developed, seeds. We put a mix of 20 seeds from one population from different mother plants into one Petri dish (5 cm in diameter) with three layers of filter paper. In each case, we created 20 Petri dishes composed of seeds from exactly the same maternal plants in the same proportions. These identical 20 dishes were divided between 4 treatments – 5 for each treatment (warm/cold × wet/dry). Another mixture of mother plants was created for other 20 Petri dishes of seeds from one population. This means that we had 2 sets of dishes (40 dishes in total) from each population. For ALP3, ALP4, SUB1 and SUB4 we were able to create only 10 identical Petri dishes and 2 different sets of genotype mixtures (20 dishes in total from one population), due to shortage of seeds. These populations were exposed to warm-wet and cold-wet regimes only, the dry regimes were not established for these.

We used 4 germination conditions in growth chambers, further referred to as target conditions. Specifically, we combined 2 levels of temperature × 2 levels of moisture. Two temperature conditions, warm and cold, were derived from long term measurements at the localities in Norway (for details see (Tingstad et al. 2015)) and were based on temperatures of the four warmest months of the year. The day temperature corresponded to average highest day temperatures at the coldest and the warmest localities of origin of the plants, the night temperature corresponded to the mean of the lowest night temperatures at the same localities. The resulting night/day temperature was 3°C/12.5°C (as a proxy for alpine conditions). The warm condition was set to 3°C/24.5°C (as a proxy for boreal conditions). The daylight period lasted from 6-22 o’clock. Dawn and twilight lasted for 2 hours before and after the night. The night period lasted from 24 till 4 o’clock.

The wet conditions were simulated by watering with demineralized water (water potential 0.0 MPa) and dry conditions by decreased water potential by addition of polyethylene glycol (PEG, molecular weight 6000) into the water. The water potential was kept approximately −0.7 MPa (intermediate dry) throughout the experiment. According to (Young and Nobel 1986), this water potential approximately corresponds to the rainfall of 600 mm. According to (Evans and Etherington 1990), such water potential represents intermediately dry conditions. The solutions of PEG were prepared and adapted to each germination temperature according to (Michel 1983). PEG adjust water potential without affecting seed germination in other way than due to the moisture itself (Hardegree and Emmerich 1994). From the 12^th^ week of the experiment, we were using more water in the PEG solution (40% more water i.e. approximately −0.4MPa), because a lot of dishes did not germinate at all.

The germinated seeds were recorded every week. The seed was considered germinated if the radicle was visible to the naked eye. Positions of Petri dishes were shifted randomly after a week. Mouldy seeds with decomposed embryo were removed from the dish. When at least 60% of the seeds had germinated and no further seeds germinated for two subsequent weeks in a specific germination conditions, we applied Gibberellic acid in concentration 0.05 g/100 ml of demineralized water to the seeds to stimulate germination (Kahn 1960). After Gibberellic acid application, we continued seed germination recording every week. The germination after application of the Gibberellic acid was used to assess seed viability, but these seeds were not scored as ‘germinated’ within the main experiment. The experiment was terminated four weeks after application of the Gibberellic acid. At this time, we removed all rotten seeds from the dishes. Healthy looking ungerminated seeds were tested for viability by tetrazolium chloride method according to (Cottrell 1947). Because there were many healthy ungerminated seeds and the viability test is very time consuming, we only selected subsets from each population and treatment. In this way, we proved that the healthy-looking ungerminated seeds can really be considered as viable.

### Data analysis

#### Effects of seed origin on nutrients and seed mass

Pair-wise correlation matrix, based on Pearson’s correlation coefficient, of all studied nutrients (N, P, starch and fructans) and seed mass is presented in Supporting information 1. Because of strong correlation of all nutrients with seed mass (all r ≥ |0.427|), we used only seed mass for testing the effect on germination behavior. To observe if the negative relationship between nitrogen resp. phosphorus and seed mass is caused by dilution of the content in case of larger seeds, we multiplied concentration of N, P, starch and fructans by seed mass. We refer to these values as contents of nutrients. We compared their relationship with seed mass (Supporting information 1).

We used ANOVA to test effect of original temperature and moisture and of mother plant nested within population on seed mass and all nutrients (N, P, starch and fructans, both concentration and content). As results on nutrient content were similar to results of nutrient concentration, we present them only in Supporting information 2.

#### Germination patterns

Germination behavior was defined as total germination, the time to 50% germination (T50), germination index (GI – describes ratio of the germination percentage and speed), proportion of dormant seeds and seed viability. Total germination was defined as the sum of germinated seeds in one Petri dish over the period before application of the Gibberellic acid. Proportion of dormant seeds was defined as the proportion of all seeds that germinated only after Gibberellic acid application or were found to be viable after tetrazolium chloride application. Proportion of viable seeds was defined as sum of germinated seeds during whole experiment (before and after Gibberellic acid application) and seeds found to be viable after tetrazolium chloride application. GI was calculated with formula from (Liu et al. 2014):

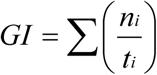

where *n*_*i*_ is the cumulative number of germinated seeds in time *t*_*i*_ (in our case one-week intervals)

T_50_ was calculated by following formula:

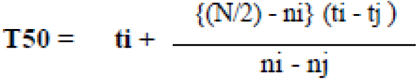

N is the final number of germinated seeds and n_i_ and n_j_ are cumulative number of seeds germinated by adjacent counts at times (weeks) t_i_ and t_j_ when n_i_<N/2< n_j_ (Sadeghi et al. 2011). GI express the initial slope of the germination curve - greater GI indicates faster initial germination. T50 covers the complete germination curve. Lower T50 means overall faster germination. In cases, when no seeds germinated the value for total germination was zero and the value for GI and T50 was classified as a missing value. Pair-wise correlation matrix of the variables, based on Pearson’s correlation coefficient, is presented in Supporting information 2. Because of strong correlation of proportion of dormant seeds with all variables (all r ≥ 0.419) and strong correlation of GI with germination percentage (r = −0.861), we did not use proportion of dormant seeds and GI in the subsequent tests.

We tested the effect of the temperature and moisture of the origin, the target temperature and moisture, seed mass and all their interactions on germination. A significant effect of target conditions will indicate phenotypic plasticity, a significant effect of origin will indicate genetic differentiation, and the interaction between target and origin will indicate genetic differentiation in plasticity.

Due to unbalanced design caused by not including four populations into dry target conditions (for details see section Germination tests) it was not possible to reliably analyze the whole dataset. Thus, we split the dataset and did three independent sets of tests.

First set of tests was carried out for all populations (11), but only wet target conditions were included into the test. To study effect of dry target condition, the second set of tests used both wet and dry target conditions, but only for populations BOR1 to BOR4 (4 populations, highlighted by square in Fig. 1), which did not allow to study effect of original temperature. The third set of tests was done using populations BOR2, BOR3, SUB2 and SUB3 (4 populations, underlined in Fig. 1), which allows us to test effects of original temperature, moisture and target temperature and moisture. All tests were done with and without including seed mass into the statistical model. As the first of tests set with all populations without wet target conditions is the most suitable for study effect of original conditions, we present them (both with and without seed mass) in the main text. The second and the third set of tests are presented in Supporting information 3, only the effect of target moisture is described in the main text. Because germination tests were done as a mixture of mother plants in one Petri dishes (see section Germination tests), we calculated mean of seed mass for each population for these analyses.

All the tests of germination characteristics were done using general mixed effect models as implemented in the lme4 package in R (Bates et al. 2015) with population as a random factor. We assumed binomial distribution of total germination and proportion of viability seeds (information on number of germinating seeds/non-germinating and germinating + dormant/died seeds linked using c-bind function in R). T50 followed Gaussian distribution after log-transformation. All statistical analyses were done in the R programme (version 3.6.0.).

#### Correlation of plant performance and germination

To assess how the germination behavior correlates with plant performance, we created pair-wise correlation matrix based on Pearson’s correlation coefficient (Table 1) with using data describe above and the data previously published in (Münzbergová et al. 2017). For this purpose, was used whole dataset of germination data in wet regime. Single values represented means for each population in each target conditions. Both these data sets used plants originated from the same localities and the temperature and moisture settings in the growth chambers were the same as well. As vegetative plant performance is the best characterized by number of ramets, weight of aboveground biomass and weight of belowground biomass, we used these variables.

**Table 1.** Pair-wise correlation matrix of germination response and plant performance (the means across the target conditions). Significant values (≤ 0.05) of Pearson correlation coefficients are in bold.

|  | Total<br>germination | T50 | Prop.<br>viable<br>seeds |
| --- | --- | --- | --- |
| Above.biom | **0.401** | -0.216 | -0.231 |
| Below.ground | **-0.743** | **0.655** | 0.283 |
| No.ramets | **-0.435** | 0.224 | **0.521** |

Because the previous study (Münzbergová et al. 2017)providing data on plant performance was based on both wet and dry regimes, we had to re-analyse the data using only the wet regimes (identical as for germination). To do this, we used the same model as is describe in Test 1 for seed germination. As a dependent variable we used number of ramets, weight of aboveground biomass and weight of belowground biomass. All variables follow Gaussian distribution without any transformation.

## Results

### Effects of seed origin on nutrients and seed mass

We did not find any differences between mother plants in the concentration of nitrogen, phosphorus, starch and fructans (all p-values ≥ 0.259). Populations significantly differed in the concentration of nitrogen (F = 3.37, p = 0.033), with the highest concentration in seed originated from cold, wet conditions (ALP3) and the lowest concentration in seed originated from warm, wet conditions (BOR3). We found marginally significant difference in concentration of phosphorus (F = 2.76, P = 0.061), with the highest content in cold, wet conditions (ALP3) and the lowest content in cold, dry conditions (ALP1). Concentrations of starch and fructans did not differ between population (F = 1.95, p = 0.143 resp. F = 0.90, p = 0.454). Effect of original conditions on content of nutrients were similar to results of nutrient concentration (Supporting information 2).

Seed mass negatively correlated with concentration of nitrogen and phosphorus and positively with starch and fructans (Supporting information 1). The relationship between nitrogen resp. phosphorus and seed mass disappear after multiplying their concentration by seed mass, i.e. calculating their total content (Supporting information 1). The highest seed mass was observed in the seeds coming from the warm and wet conditions and the lowest in the seeds from cold and wet conditions.

### Effect of seed mass, original and target conditions on germination

Test including also seed mass as a predictor showed that target temperature significantly affected all germination variables (Table 2). Total germination and proportion of viable seeds increased and T50 decreased in warmer target temperature. Original temperature significantly influenced total germination and proportion of viable seeds (Table 2) with higher total germination and proportion of viable seeds in seeds coming from warm conditions. Original moisture significantly affected only T50 (Table 2) with the fastest germination in seeds coming from the wettest condition and the slowest germination in seeds coming from the driest conditions. Seed mass significantly affected all the study variables (Table 2). Total germination and proportion of viable seeds increased and T50 decreased with increasing seed mass.

**Table 2.** Effect of seed mass, original (O) and target (T) conditions on total germination, proportion of viable seeds and time to 50% germination (T50) assessed using mixed effect models with population used as a random factor. Table represents results all populations without wet regime (the first analysis). Significant values (≤ 0.05) are in bold. * indicate significant result in the model not including seed mass. ● indicate non-significant results in the model not including seed mass.

|  | Total germination |  | Prop. viable seeds |  | T50 |  |
| --- | --- | --- | --- | --- | --- | --- |
|  | F-values | p-value | F-values | p-value | F-values | p-value |
| TTemp | **32.27** | **<0.001** | **23.53** | **<0.001** | **6.85** | **0.010** |
| OTemp | **6.60** | **<0.001•** | **3.66** | **0.038•** | 1.29 | 0.265 |
| OMoist | 1.70 | 0.152 | 1.07 | 0.304 | **4.25** | **0.048•** |
| Seed mass | **19.39** | **<0.001** | **13.49** | **<0.001** | **9.44** | **0.004** |
| TTemp: OTemp | 0.41 | 0.351 | 1.00 | 0.356 | 0.88 | 0.350 |
| TTemp: OMoist | 0.13 | 0.700 | 0.14 | 0.693 | 2.92 | 0.089 |
| OTemp: OMoist | 0.13 | 0.701* | 0.05 | 0.738 | 0.28 | 0.600 |
| TTemp: Seed mass | **4.05** | **0.017** | **5.03** | **0.009** | **5.48** | **0.020** |
| OTemp: Seed mass | **4.59** | **0.019** | **2.67** | **0.036** | **4.98** | **0.033** |
| OMosit: Seed mass | 1.38 | 0.194 | 0.84 | 0.396 | 3.90 | 0.057 |
| TTemp: OTemp: OMoist | 0.24 | 0.656 | 0.50 | 0.637 | 0.30 | 0.586 |
| TTemp: OTemp: Seed mass | 0.21 | 0.598 | 0.00 | 0.859 | 2.93 | 0.089 |
| TTemp: OMoist: Seed mass | 0.06 | 0.741 | 0.11 | 0.751 | 2.32 | 0.130 |
| OTemp: OMoist: Seed mass | 0.80 | 0.438 | 1.39 | 0.215 | 1.91 | 0.177 |

We found only two significant double interaction, specifically interaction of target temperature with seed mass and original temperature with seed mass (Table 2). Interaction of target temperature and seed mass significantly influenced all study variables (Table 2). Total germination and proportion of viable seeds were the highest in heavy seeds exposed to warm target temperature and the lowest in light seeds exposed to cold target temperature (Fig. 2). T50 was the highest (i.e. germination speed was low) in light seeds exposed to cold target temperature and the lowest (i.e. germination speed was high) in heavy seeds exposed to warm target temperature. In warm target temperature the differences between different seed mass was lower than in cold target temperature (Fig. 2). Interaction of original temperature and seed mass significantly influenced all study variables (Table 2). Total germination and proportion of viable seed were the highest in heavy seeds coming from the warmest conditions and the lowest in light seeds coming from colder conditions (Fig. 3). T50 was the highest (i.e. germination speed was low) in light seeds coming from the warmest conditions and the lowest (i.e. germination speed was high) in heavy seeds coming from the warm conditions (Fig. 3).

**Fig 2.**
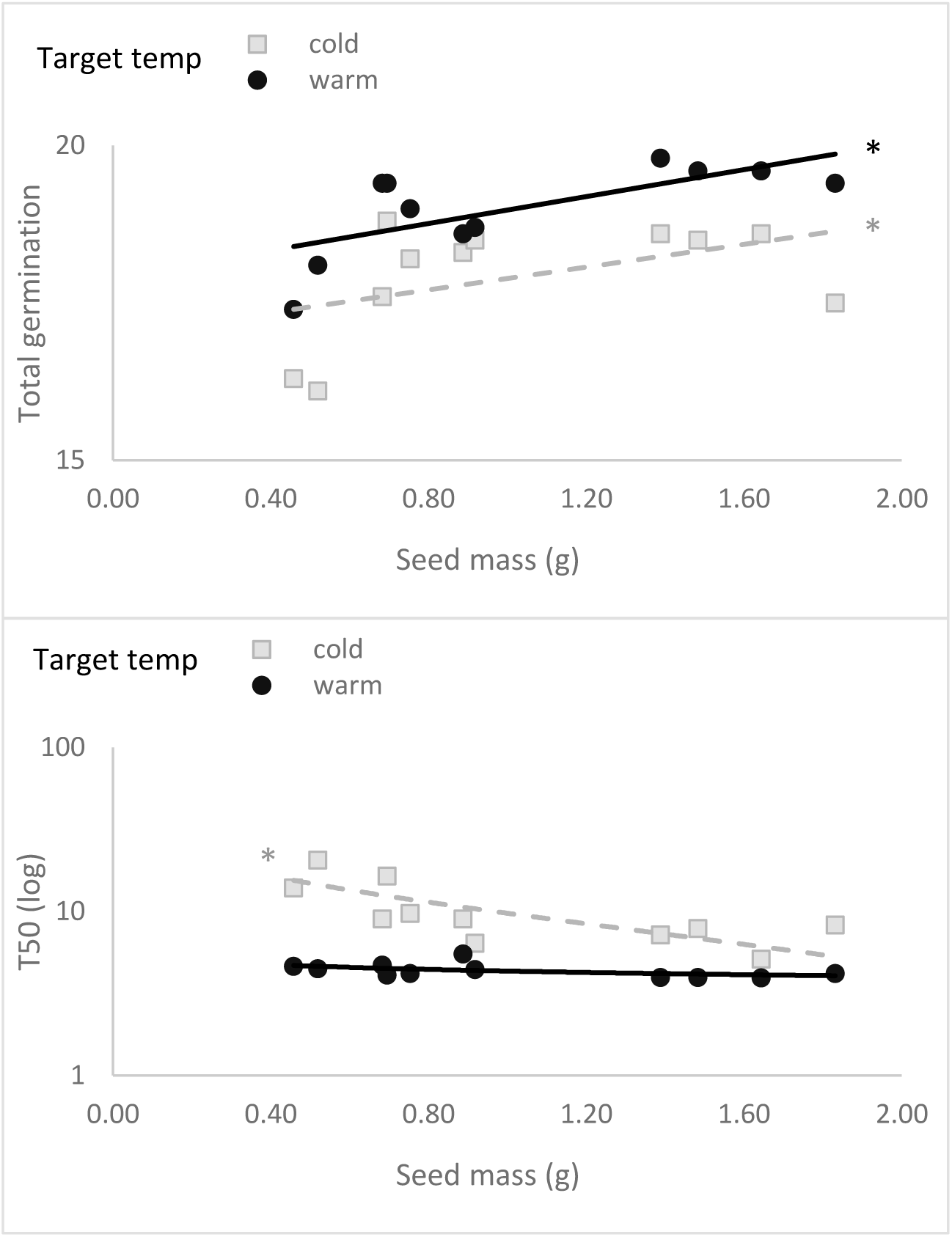
Effect of seed mass and target temperature on A) total germination (proportion) and B) T50. In both cases, there is a significant interaction between target temperature and seed mass. The graphs show means of total germination resp. T50. * significantly different

**Fig 3.**
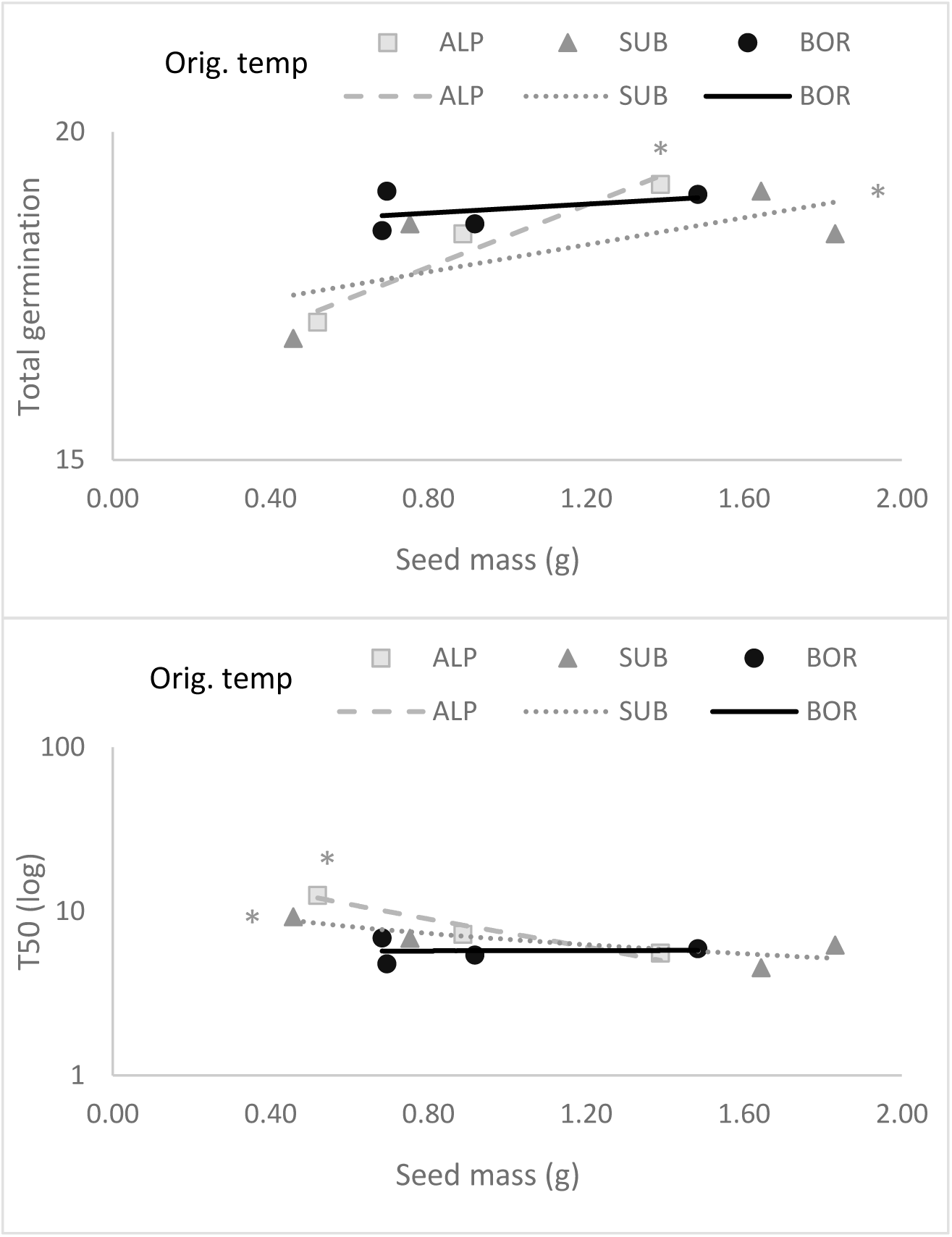
Effect of seed mass and original temperature on A) total germination (proportion) and B) T50. In both cases, there is a significant interaction between target temperature and seed mass. The graphs show means of total germination resp. T50. * significantly different

The results of the model not including seed mass were largely the same as above with few exceptions. Effect of original temperature and moisture became non-significant when seed mass was not included in the model (Table 2). In contrast, we found significant effect of interaction of original moisture and temperature on total germination in this case (Table 2). The highest total germination was in seeds coming from the warmest and wettest conditions and the lowest in seeds coming from the coldest and intermediate wet conditions.

The other two models, which included target moisture, showed significant effect of target moisture on all study variables (Supporting information 3), with higher total germination in wet condition and higher proportion viable seeds and T50 in dry conditions. The most interesting interactions were the following. Target moisture significantly interacted with target temperature and affected total germination and proportion of viable seeds (Supporting information 3). The highest values of total germination were in wet and warm conditions and the lowest in dry and cold conditions. Proportion of viable seeds had the lowest numbers in wet and cold target conditions and the highest numbers in dry and cold target conditions. Target moisture interacted with original temperature and affected proportion of viable seeds (Supporting information 3), with the highest numbers in seeds coming from warmer condition and exposed to dry target condition. The lowest numbers were observed in seeds coming from colder conditions and exposed to wet target conditions. In these two models were detected interactions of target and original conditions (Supporting information 3).

### Correlation of plant performance and germination response

Total germination positively correlated with aboveground biomass and negatively with belowground biomass and number of ramets (Table 1). T50 positively correlated only with belowground biomass and number of viability seeds positively correlated with number of ramets (Table 1). No other correlations between plant performances and germination responses were significant.

Target temperature significantly affected all germination characteristics, as described above, as well as belowground biomass, aboveground biomass and marginally number of ramets (F = 5.25, p = 0.022, resp. F = 7.59, p = 0.006, resp. F = 2.98, p = 0.085). Original temperature and moisture had significant effect on germination (Table 2), but not on any of the vegetative characteristics (p ≥ 0.248). In contrast, belowground biomass (F = 5.05, p = 0.025) and number of ramets (F = 3.19, p = 0.075, marg. sig.) were affected by interaction of target temperature and original moisture, while germination characteristics were not (Table 2). We did not find any other significant effects of interactions on germination (Table 2, except interactions with seed mass) or on plant performance (p ≥ 0.096).

## Discussion

Seed mass, concentration of nitrogen and marginally phosphorus, but not of carbohydrates were affected by original conditions. Seed mass correlated with all the study nutrients with heavier seeds having higher concentration of carbohydrates and lower concentration of nitrogen and phosphorus, which is caused by dilution the same amout of nitrogen and phosphorus in larger seeds. In all analyses, increasing seed mass had positive impact on germination (higher and faster germination), while effect of original conditions was significant only in some analyses. Significant results show higher germination and seed viability in seeds coming from the warmest condition. Warm target temperature increased germination, proportion of viable seeds and decreased T50. Target temperature and moisture interacted, but original and target conditions not. Wet and warm target conditions were suitable for germination of all populations. In other germination conditions, which were less suitable for germination, we found high proportion of viable seeds. Total germination correlated with all plant performance characteristics, with strong negative correlation with belowground biomass and number of ramets and positive correlation with aboveground biomass.

### Effects of seed origin on nutrients and seed mass

Higher seed mass is commonly expected to be associated with higher concentration of seed nutrients (Wulff and Bazzaz 1992) (Rees et al. 2001). Intraspecifically, few studies confirm this for nitrogen, phosphorus and sulfur (Vaughton and Ramsey 2001) (Obeso 2012) (Kolodziejek 2017). In our study, all nutrients are correlated with seed mass, however, only starch and fructans positively. As expected, only carbohydrates contributed positively to seed mass as they have higher molecular weight than nitrogen and phosphorus. Concentration of nitrogen and phosphorus negatively correlated with seed mass. Similar result for phosphorus was found in (Mašková and Herben in prep) at the interspecific level. The possible explanation of our results is that the same amount of nitrogen and phosphorus is more diluted in larger seeds and it was confirmed as relationship became non-significant for nutrient content.

We found significant differences between populations both in the concentration and content of nitrogen with the highest values in seed coming from the coldest and the wettest conditions and marginally significant differences between populations in the concentration of phosphorus with the same trend. Possible explanation could be that plants originated from the coldest conditions in Norway made low quality seeds in common garden conditions that in their home conditions. All plants performed well in the experimental garden, and it is thus unlikely that quality of the seeds would be affected by low performance of some of the plants in the garden and at interspecific level (Bu et al. 2018) demonstrated higher concentration of nitrogen and phosphorus in the highest altitude i.e. in the coldest and the wettest conditions as well. It seems that these conditions are not the most suitable for seed recruitment and species invest the energy to these specific seed nutrients to supported seed establishment before developed root system. The higher nitrogen and phosphorus concentration of seeds from specific conditions might derive from adaptive selection (Bu et al. 2018). We did not find any differences between mother plants within populations, which support the claim about adaptive selection. Only those mother plants, which sufficiently invest to seed nutrients grow on study localities and simultaneously they do not invest more than it is necessary for their specific locality.

Differences in concentration of nutrients in seeds could be caused by soil compositions (Lukac et al. 2010). It was demonstrated that plant growing in poor soil have higher concentration of seed nutrients (Mašková and Herben in prep). Our study localities were chosen to ensure that bedrock was as similar as possible. We thus expect that effect of soil composition is much less important than the climate itself.

We observed differences between populations in seed mass, which could be attributed to original climatic conditions (Winn and Gross 1993) (Zas and Sampedro 2015) (Gorden et al. 2016), with higher seed mass in warmer (Gorden et al. 2016) and wetter conditions (Wulff and Bazzaz 1992) (Gorden et al. 2016), which is in line with our results. It is possible that ongoing changes in temperature and precipitation will have impact on seed mass, which could further affect for instance dispersal, predation, seed-bank persistent or seedling establishment.

### Effect of seed mass, original and target conditions on germination

Many previous studies suggested that larger seeds have higher and faster germination (e.g. (Greipsson and Davy 1995) (Münzbergová and Plačková 2010) (Wu and Du 2007) (Paulů et al. 2017) (Veselá et al. 2019)). This is commonly attributed to higher concentration of nutrients, which nourish the sprout (Rees et al. 2001). Our study demonstrated that higher seed mass is caused by higher concentration only of carbohydrates not by nitrogen and phosphorus. Therefore, higher concentration of carbohydrates could be contributed to higher and faster germination, however, more extensive studies are required to confirm this.

Seed mass interacted with target temperature, with the fastest and highest germination in heavy seeds exposed to warm condition. This result could indicate that carbohydrates are more effectively used in warm target conditions. In our previous study, we showed, that seed mass interacted with original conditions with higher and faster germination in seed coming from warm and dry localities (Veselá et al. 2019). In the current study, we observed a similar trend but only for the interaction of seed mass with original temperature and not moisture. To our best knowledge no other study showed the interaction of seed mass with original climatic conditions.

Original temperature itself significantly affected total germination and proportion of viable seeds. In line with our expectation, higher values were observed in seeds coming from warm conditions. This is in line with other studies (Cruz et al. 2003) (Ndihokubwayo et al. 2016) (Santo et al. 2015) (Bauk et al. 2017) (Mira et al. 2017) (Veselá et al. submitted). This result could be caused by longer vegetation period in lowlands allowing seeds to properly develop (Milbau et al. 2009) (Meineri et al. 2013). It could also be explained by higher seed mass in seed originated from warmer conditions as described above. Germination response of *F. rubra* was affected by original moisture, but only for T50. Similar result was found in (Veselá et al. submitted^1^). However, both the effects of original temperature and moisture were significant only in the model including seed mass. This result could indicate that seed mass masks the effect of original conditions.

It is known that target conditions have strong impact on species germination (e.g. (Grime et al. 1981, Schütz and Rave 1999) (Gardarin et al. 2011)). Our results are in line with other studies demonstrating that germination increases in higher target temperatures (Gardarin et al. 2011) (Ooi, Auld and Denham 2012) (Walder and Erschbamer 2015), but see (Veselá et al. 2019). Temperature and moisture can interact and significantly influence germination response (Rivas-Arancibia et al. 2006) (Gurvich et al. 2017) (Veselá et al. submitted) as in the case of *F. rubra*. Total germination and T50 was negatively affected by decreasing target moisture both in cold and warm target temperature (similarly e.g. (Wen et al. 2015) (Ruhl et al. 2015)), but high proportion of viable seeds in all dry target conditions indicate seed ability to survive period of drought and germinate in wet season. Significant effects of target conditions indicate high variability of germination response of *F. rubra*, which may enable species to tolerate fluctuating climate conditions (Wainwright and Cleland 2013).

Interaction of original and target climate indicates genetic differentiation in plasticity (*sensu* Pigliucci 2001). Only a few studies focused on this in germination (reviewed in (Walck et al. 2011). We expected that original and target conditions will interact similarly like in our study (Veselá et al. submitted), but no such interactions have been detected in the main results of this study. This could be caused we did not include target moisture into the main analysis. These interactions were found in additional analyses studying target moisture.

Phenotypic plasticity can be defined as ‘the ability of one genotype to produce different phenotypes when exposed to different environmental conditions’ (Pigliucci, Murren and Schlichting 2006). It follows that cases, when seeds do not germinate and died, cannot be considered as a phenotypic plasticity. Contrariwise cases, when seeds do not germinate and stay dormant, can be use term phenotypic plasticity, as dormancy is response on environmental condition similarly as germination speed (Baskin and Baskin 2001).

### Correlation of plant performance and germination response

The investment of clonal plants into either vegetative spread or generative reproduction is usually modified by the environment, with vegetative spread being favoured in less competitive environment (Eriksson 1992) (Chaloupecka and Leps 2004), where species have sufficient sources for building biomass and high-quality seeds.

Many studies have detected negative correlation between generative and vegetative reproduction (e.g. (Cheplick 1995) (Worley and Harder 1996) (Ronsheim and Bever 2000) (van Kleunen et al. 2002) (Herben et al. 2012)), while others have not (e.g. (Reekie 1991) (Cain and Damman 1997)). Negative correlations of total germination with belowground biomass and number of ramets confirm negative correlation between generative and vegetative reproduction. Number of ramets positively correlated with proportion of viable seeds, which is in line with previous correlation, as proportion of viable seeds include both germinated and dormant seeds. It is highly probable that in cases of high proportion of dormant seeds, growth of ramets is supported. Positive correlation of belowground biomass and T50 support the statement described above, since with slow germination (high T50), production of vegetative organs is preferred. All the results indicate that *F. rubra* shows trade-off between generative and vegetative reproduction.

It was demonstrated that target conditions affect plant performance (Münzbergová et al. 2017) as well as germination (Gardarin et al. 2011) (Walder and Erschbamer 2015). We show that target temperature affected plant performance and germination, in plants originated from the same localities and exposed to the same target temperatures. This result indicates that *F. rubra* has high phenotypic plasticity both for plant performance and germination and species will probably successfully cope with ongoing climate change. We did not find effect of original climate on plant performance, despite the previous study providing plant performance data (Münzbergová et al. 2017) found this effect, which indicate absence of genetic differences. This may be because our study used only plants cultivated under wet target conditions, while (Münzbergová et al. 2017) used complete dataset. This result suggests that genetic differences in plant performance may be detected just under some conditions. While drier conditions seem as more stressful for plant germination, these same conditions are probably less stressful for plant growth (as the plants in wet conditions suffered from water logging).

## Conclusion

To our best knowledge, this is the first study demonstrating relationship between seed mass and specific seed nutrients across localities experiencing different climatic conditions. Our results indicate that higher seed mass is caused by higher concentration of carbohydrates. Further, it seems that higher seed mass, and probably higher concentration of carbohydrates, may play role in higher and faster germination in seeds coming from warmer localities and simultaneously also in higher and faster germination in warm target conditions. With regards that seeds mass is strongly affected by climate, it is possible that ongoing climate change will have impact on seed mass, which could further affect for instance dispersal, predation, seed-bank persistent or seedling establishment. Germination response of *F. rubra*, as a dominant species of meadows, shows high proportion of germinated or dormant seeds, which suggest phenotypic plasticity of study species. Phenotypic plasticity allows to species cope with changing climate. When the target conditions were unsuitable and germination was low, seeds stay dormant and probably they would be able to germinate in more suitable condition. Simultaneously, *F. rubra* show trade-off between generative and vegetative reproduction indicating that the species is able to modify investments to generative or vegetative reproduction depending on the actual conditions. The importance of target and origin for germination cannot be, however, easily predicted from their effects on growth. Both these types of variables should thus be studied in future studies to obtain an overall picture of plant performance under changing climates.

## Acknowledgement

We thank J. Knappová and V. Hadincová for help with collection of the maternal plants in the field and V. Hadincová, M. Lokvencová, I. Jarošincová and I. Chmelařová for help with plant cultivation and seed collection in the experimental garden and participants of POPECOL seminars on constructive comments on the previous version of the manuscript. The study was supported by project GAČR 19-00522S.

## References

Anderson, J. T. & Z. J. Gezon (2015) Plasticity in functional traits in the context of climate change: a case study of the subalpine forb Boechera stricta (Brassicaceae). Global Change Biology, 21, 1689–1703.

Anderson, J. T., D. W. Inouye, A. M. McKinney, R. I. Colautti & T. Mitchell-Olds (2012) Phenotypic plasticity and adaptive evolution contribute to advancing flowering phenology in response to climate change. Proceedings of the Royal Society B-Biological Sciences, 279, 3843–3852.

Bates, D., M. Machler, B. M. Bolker & S. C. Walker (2015) Fitting Linear Mixed-Effects Models Using lme4. Journal of Statistical Software, 67, 1–48.

Bauk, K., J. Flores, C. Ferrero, R. Perez-Sanchez, M. L. L. Penas & D. E. Gurvich (2017) Germination characteristics of Gymnocalycium monvillei (Cactaceae) along its entire altitudinal range. Botany, 95, 419–428.

Bu, H. Y., P. Jia, W. Qi, K. Liu, D. H. Xu, W. J. Ge & X. J. Wang (2018) The effects of phylogeny, life-history traits and altitude on the carbon, nitrogen, and phosphorus contents of seeds across 203 species from an alpine meadow. Plant Ecology, 219, 737–748.

Cain, M. L. & H. Damman (1997) Clonal growth and ramet performance in the woodland herb, Asarum canadense. Journal of Ecology, 85, 883–897.

Chaloupecka, E. & J. Leps (2004) Equivalence of competitor effects and tradeoff between vegetative multiplication and generative reproduction: case study with Lychnis flos-cuculi and Myosotis nemorosa. Flora, 199, 157–167.

Cheplick, G. P. (1995) Life-history trade-offs in *Amphibromus scabrivalies* (Poaceae) - allocation to clonal growth, storage, adn cleistoamous reproduction. American Journal of Botany, 82, 621–629.

Cottrell, H. J. (1947) Tetrazolium salt as a seed germination indicator. Nature, 159, 748–748.

Cruz, A., B. Perez, A. Velasco & J. M. Moreno (2003) Variability in seed germination at the interpopulation, intrapopulation and intraindividual levels of the shrub Erica australis in response to fire-related cues. Plant Ecology, 169, 93–103.

Dalgleish, H. J., D. N. Koons & P. B. Adler (2010) Can life-history traits predict the response of forb populations to changes in climate variability? Journal of Ecology, 98, 209–217.

Degreef, J., O. J. Rocha, T. Vanderborght & J. P. Baudoin (2002) Soil seed bank and seed dormancy in wild populations of lima bean (Fabaceae): Considerations for in situ and ex situ conservation. American Journal of Botany, 89, 1644–1650.

Eriksson, O. (1992) Evolution of seed dispersal and recruitment in clonal plants. Oikos, 63, 439–448.

Evans, C. E. & J. R. Etherington (1990) The effect of soil-water potential on seed-germination of some British plants. New Phytologist, 115, 539–548.

Fay, P. A. & M. J. Schultz (2009) Germination, survival, and growth of grass and forb seedlings: Effects of soil moisture variability. Acta Oecologica-International Journal of Ecology, 35, 679–684.

Gardarin, A., C. Daurr & N. Colbach (2011) Prediction of germination rates of weed species: Relationships between germination speed parameters and species traits. Ecological Modelling, 222, 626–636.

Gorden, N. L. S., K. J. Winkler, M. R. Jahnke, E. Marshall, J. Horky, C. Huddelson & J. R. Etterson (2016) Geographic patterns of seed mass are associated with climate factors, but relationships vary between species. American Journal of Botany, 103, 60–72.

Greipsson, S. & A. J. Davy (1995) Seed mass and germination behavior in populations of the denu-duildinf grass Leymus arenarius. Annals of Botany, 76, 493–501.

Grime, J. P., G. Mason, A. V. Curtis, J. Rodman, S. R. Band, M. A. G. Mowforth, A. M. Neal & S. Shaw (1981) A comparative study of germination characteristics in a local flora. Journal of Ecology, 69, 1017–1059.

Gurvich, D. E., R. Perez-Sanchez, K. Bauk, E. Jurado, M. C. Ferrero, G. Funes & J. Flores (2017) Combined effect of water potential and temperature on seed germination and seedling development of cacti from a mesic Argentine ecosystem. Flora, 227, 18–24.

Hardegree, S. P. & W. E. Emmerich (1994) Seed-germination response to polyethylene-glycol solution depth. Seed Science and Technology, 22, 1–7.

Herben, T., F. Krahulec, V. Hadincova & S. Pechackova (2001) Clone-specific response of Festuca rubra to natural variation in biomass and species composition of neighbours. Oikos, 95, 43–52.

Herben, T., Z. Novakova, J. Klimesova & L. Hrouda (2012) Species traits and plant performance: functional trade-offs in a large set of species in a botanical garden. Journal of Ecology, 100, 1522–1533.

Huntley, B. (1991) How plants respond to climate change - migration rates, individualism and the consequences for plant-communities. Annals of Botany, 67, 15–22.

Jump, A. S. & J. Penuelas (2005) Running to stand still: adaptation and the response of plants to rapid climate change. Ecology Letters, 8, 1010–1020.

Kahn, A. (1960) Promotion of lettuce seed germination by gibberellin. Plant Physiology, 35, 333–339.

Klanderud, K., V. Vandvik & D. Goldberg (2015) The Importance of Biotic vs. Abiotic Drivers of Local Plant Community Composition Along Regional Bioclimatic Gradients. Plos One, 10, e0130205.

Knappová, J., D. Židlická, T. Kadlec, M. Knapp, D. Haisel, V. Hadincová & Z. Münzbergová (2018) Population differentiation related to climate of origin affects the intensity of plant-herbivore interactions in a clonal grass. Basic and Applied Ecology, 28, 76–86.

Kolb, A., J. P. Dahlgren & J. Ehrlen (2010) Population size affects vital rates but not population growth rate of a perennial plant. Ecology, 91, 3210–3217.

Kolodziejek, J. (2017) Effect of seed position and soil nutrients on seed mass, germination and seedling growth in Peucedanum oreoselinum (Apiaceae). Scientific Reports, 7.

Laughlin, D. C., R. T. Strahan, P. B. Adler & M. M. Moore (2018) Survival rates indicate that correlations between community-weighted mean traits and environments can be unreliable estimates of the adaptive value of traits. Ecology Letters, 21, 411–421.

Lloret, F., J. Penuelas & M. Estiarte (2004) Experimental evidence of reduced diversity of seedlings due to climate modification in a Mediterranean-type community. Global Change Biology, 10, 248–258.

Lukac, M., C. Calfapietra, A. Lagomarsino & F. Loreto (2010) Global climate change and tree nutrition: effects of elevated CO2 and temperature. Tree Physiology, 30, 1209–1220.

McCleary, B. V., T. S. Gibson, V. Solah & D. C. Mugford (1994) Total starch measurement in cereal products - interlaboratory evaluation of a rapid enzymatic test procedure. Cereal Chemistry, 71, 501–505.

McGinley, M. A. & E. L. Charnov (1988) Multiple resources and the optimal balance between size and number of offspring. Evolutionary Ecology, 2, 77–84.

Meineri, E., O. Skarpaas, J. Spindelbock, T. Bargmann & V. Vandvik (2014) Direct and size-dependent effects of climate on flowering performance in alpine and lowland herbaceous species. Journal of Vegetation Science, 25, 275–286.

Meineri, E., J. Spindelbock & V. Vandvik (2013) Seedling emergence responds to both seed source and recruitment site climates: a climate change experiment combining transplant and gradient approaches. Plant Ecology, 214, 607–619.

Meyer, S. E., P. S. Allen & J. Beckstead (1997) Seed germination regulation in *Bromus tectorum* (Poaceae) and its ecological significance. Oikos, 78, 475–485.

Michel, B. E. (1983) Evaluation of the water potentials of solutions of polyethylene glycol-8000 both in the absence and presence of other solutes. Plant Physiology, 72, 66–70.

Milbau, A., B. J. Graae, A. Shevtsova & I. Nijs (2009) Effects of a warmer climate on seed germination in the subarctic. Annals of Botany, 104, 287–296.

Mira, S., A. Arnal & F. Perez-Garcia (2017) Habitat-correlated seed germination and morphology in populations of Phillyrea angustifolia L. (Oleaceae). Seed Science Research, 27, 50–60.

Münzbergová, Z. (2005) Determinants of species rarity: Population growth rates of species sharing the same habitat. American Journal of Botany, 92, 1987–1994.

Münzbergová, Z. & V. Hadincová (2017) Transgenerational plasticity as an important mechanism affecting response of clonal species to changing climate. Ecology and Evolution, ECE33105.

Münzbergová, Z., V. Hadincová, H. Skálová & V. Vandvik (2017) Genetic differentiation and plasticity interact along temperature and precipitation gradients to determine plant performance under climate change. Journal of Ecology, DOI: 10.1111/1365-2745.12762.

Münzbergová, Z., V. Latzel, M. Šurinová & V. Hadincová (2018) DNA methylation as a possible mechanism affecting ability of natural populations to adapt to changing climate. Oikos.

Münzbergová, Z. & I. Plačková (2010) Seed mass and population characteristics interact to determine performance of Scorzonera hispanica under common garden conditions. Flora, 205, 552–559.

Ndihokubwayo, N., V. T. Nguyen & D. D. Cheng (2016) Effects of origin, seasons and storage under different temperatures on germination of Senecio vulgaris (Asteraceae) seeds. Peerj, 4.

Neilson, R. P., L. F. Pitelka, A. M. Solomon, R. Nathan, G. F. Midgley, J. M. V. Fragoso, H. Lischke & K. Thompson (2005) Forecasting regional to global plant migration in response to climate change. Bioscience, 55, 749–759.

Nicotra, A. B., O. K. Atkin, S. P. Bonser, A. M. Davidson, E. J. Finnegan, U. Mathesius, P. Poot, M. D. Purugganan, C. L. Richards, F. Valladares & M. van Kleunen (2010) Plant phenotypic plasticity in a changing climate. Trends in Plant Science, 15, 684–692.

Obeso, J. R. (2012) Mineral nutrient stoichiometric variability in Hedera helix (Araliaceae) seeds. Annals of Botany, 109, 801–806.

Ooi, M. K. J., T. D. Auld & A. J. Denham (2012) Projected soil temperature increase and seed dormancy response along an altitudinal gradient: implications for seed bank persistence under climate change. Plant and Soil, 353, 289–303.

Paulů, A., L. Harčariková & Z. Münzbergová (2017) Are there systematic differences in germination between rare and common species? A case study from central European mountains. Flora, 236-237, 15–24.

Pigliucci, M. (2001) Phenotypic Plasticity: Beyond Nature and Nurture. The Johns Hopkins University Press, Baltimore, Maryland, USA.

Pigliucci, M., C. J. Murren & C. D. Schlichting (2006) Phenotypic plasticity and evolution by genetic assimilation. Journal of Experimental Biology, 209, 2362–2367.

Qaderi, M. M. & P. B. Cavers (2002) Interpopulation and interyear variation in germination in Scotch thistle, Onopordum acanthium L., grown in a common garden: Genetics vs environment. Plant Ecology, 162, 1–8.

Reekie, E. G. (1991) Cost of seed versus rhizome production in *Agropyron repens*. Canadian Journal of Botany-Revue Canadienne De Botanique, 69, 2678–2683.

Rees, M., R. Condit, M. Crawley, S. Pacala & D. Tilman (2001) Long-term studies of vegetation dynamics. Science, 293, 650–655.

Rivas-Arancibia, S. P., C. Montana, J. X. V. Hernandez & J. A. Zavala-Hurtado (2006) Germination responses of annual plants to substrate type, rainfall, and temperature in a semi-arid intertropical region in Mexico. Journal of Arid Environments, 67, 416–427.

Ronsheim, M. L. & J. D. Bever (2000) Genetic variation and evolutionary trade-offs for sexual and asexual reproductive modes in Allium vineale (Lillaceae). American Journal of Botany, 87, 1769–1777.

Ruhl, A. T., R. L. Eckstein, A. Otte & T. W. Donath (2015) Future challenge for endangered arable weed species facing global warming: Low temperature optima and narrow moisture requirements. Biological Conservation, 182, 262–269.

Santo, A., E. Mattana & G. Baechetta (2015) Inter- and intra-specific variability in seed dormancy loss and germination requirements in the Lavatera triloba aggregate (Malvaceae). Plant Ecology and Evolution, 148, 100–110.

Schütz, W. & G. Rave (1999) The effect of cold stratification and light on the seed germination of temperate sedges (Carex) from various habitats and implications for regenerative strategies. Plant Ecology, 144, 215–230.

Skalova, H., S. Pechackova, J. Suzuki, T. Herben, T. Hara, V. Hadincova & F. Krahulec (1997) Within population genetic differentiation in traits affecting clonal growth: Festuca rubra in a mountain grassland. Journal of Evolutionary Biology, 10, 383–406.

Suseela, V., R. T. Conant, M. D. Wallenstein & J. S. Dukes (2012) Effects of soil moisture on the temperature sensitivity of heterotrophic respiration vary seasonally in an old-field climate change experiment. Global Change Biology, 18, 336–348.

Talvitie, N. A., D. P. Illustre & E. Perez (1962) Spectophotometric determination of phosphorus as molybdovanadophosphoric acid - application to air-borne particulate matter. Analytical Chemistry, 34, 866-&.

Tingstad, L., S. L. Olsen, K. Klanderud, V. Vandvik & M. Ohlson (2015) Temperature, precipitation and biotic interactions as determinants of tree seedling recruitment across the tree line ecotone. Oecologia, 179, 599–608.

Valladares, F., S. J. Wright, E. Lasso, K. Kitajima & R. W. Pearcy (2000) Plastic phenotypic response to light of 16 congeneric shrubs from a Panamanian rainforest. Ecology, 81, 1925–1936.

van Kleunen, M., M. Fischer & B. Schmid (2002) Experimental life-history evolution: Selection on the allocation to sexual reproduction and its plasticity in a clonal plant. Evolution, 56, 2168–2177.

Vaughton, G. & M. Ramsey (2001) Relationships between seed mass, seed nutrients, and seedling growth in Banksia cunninghamii (proteaceae). International Journal of Plant Sciences, 162, 599–606.

Veselá, A., T. Dostálek, M. Rokaya & Z. Münzbergová (2019) Seed mass and plant origin interact to determine species germination patterns. bioRxiv, 841114.

Wainwright, C. E. & E. E. Cleland (2013) Exotic species display greater germination plasticity and higher germination rates than native species across multiple cues. Biological Invasions, 15, 2253–2264.

Walck, J. L., S. N. Hidayati, K. W. Dixon, K. Thompson & P. Poschlod (2011) Climate change and plant regeneration from seed. Global Change Biology, 17, 2145–2161.

Walder, T. & B. Erschbamer (2015) Temperature and drought drive differences in germination responses between congeneric species along altitudinal gradients. Plant Ecology, 216, 1297–1309.

Wang, J. H., C. C. Baskin, X. L. Cui & G. Z. Du (2009) Effect of phylogeny, life history and habitat correlates on seed germination of 69 arid and semi-arid zone species from northwest China. Evolutionary Ecology, 23, 827–846.

Wen, B., P. Xue, N. Zhang, Q. Yan & M. Ji (2015) Seed germination of the invasive species Piper aduncum as influenced by high temperature and water stress. Weed Research, 55, 155–162.

Wilczek, A. M., L. T. Burghardt, A. R. Cobb, M. D. Cooper, S. M. Welch & J. Schmitt (2010) Genetic and physiological bases for phenological responses to current and predicted climates. Philosophical Transactions of the Royal Society B-Biological Sciences, 365, 3129–3147.

Winn, A. A. & K. L. Gross (1993) Latitudinal variation in seed weight and flower numbers in Prunella vulagaris. Oecologia, 93, 55–62.

Worley, A. C. & L. D. Harder (1996) Size-dependent resource allocation and costs of reproduction in Pinguicula vulgaris (Lentibulariaceae). Journal of Ecology, 84, 195–206.

Wu, G. L. & G. Z. Du (2007) Germination is related to seed mass in grasses (Poaceae) of the eastern Qinghai-Tibetan Plateau, China. Nordic Journal of Botany, 25, 361–365.

Wu, G. L., W. Li & G. Z. Du (2011) Relationship between germination and seed size in alpine shrubs in Tibetan Plateau. Pakistan Journal of Botany, 43, 2793–2796.

Wulff, R. D. & F. A. Bazzaz (1992) Effect of the parental nutrient regime on growth of the progeny in Abutilon theophrasti (Malvaceae). American Journal of Botany, 79, 1102–1107.

Young, D. R. & P. S. Nobel (1986) Predictors of soil-water potentials in the northwestern Sonoran desert. Journal of Ecology, 74, 143–154.

Zas, R. & L. Sampedro (2015) Heritability of seed weight in Maritime pine, a relevant trait in the transmission of environmental maternal effects. Heredity, 114, 116–124.

Šurinová, M., V. Hadincová, V. Vandvik & Z. Münzbergová (2019) Temperature and precipitation, but not geographic distance, explain genetic relatedness among populations in the perennial grass Festuca rubra. Journal of Plant Ecology, 12, 730–741.

