## Supplement material for "Plant origin determines seed mass, seed nutrients and germination behavior of a dominant grass species"

Supporting information 1. Pair correlation matrix, based on Pearson ´s correlation coefficient, for seed mass and A) nutrient concentration and B) nutrient content. Significant values (≤ 0.05) are in bold.

| **A** | Seed mass | Nitrogen | Phosphorus | Fructans | Starch |
|---|---|---|---|---|---|
| Seed mass | 1 | **-0.4531** | **-0.5002** | **0.6077** | **0.4265** |
| Nitrogen | | 1 | 0.0782 | **-0.6333** | **0.7195** |
| Phosphorus | | | 1 | **-0.7296** | -0.0326 |
| Fructans | | | | 1 | **-0.5943** |
| Starch | | | | | 1 |

| **B** | Seed mass | Nitrogen | Phosphorus | Fructans | Starch |
|---|---|---|---|---|---|
| Seed mass | 1 | -0.1514 | -0.2356 | **0.8480** | **0.8730** |
| Nitrogen | | 1 | 0.3386 | 0.1763 | **0.4512** |
| Phosphorus | | | 1 | 0.2472 | 0.1751 |
| Fructans | | | | 1 | 0.3216 |
| Starch | | | | | 1 |

Supporting information 2. Effect of original (O) temperature and moisture on nutrient content. Significant values (≤ 0.05) are in bold.

| | Nitrogen | | Phosphorus | | Fructans | | Starch | |
|---|---|---|---|---|---|---|---|---|
| | F-value | p-value | F-value | p-value | F-value | p-value | F-value | p-value |
| OMoist | 1.42 | 0.241 | 1.77 | 0.192 | 0.26 | 0.611 | 1.54 | 0.223 |
| OTemp | 0.01 | 0.985 | 0.98 | 0.330 | 0.04 | 0.847 | 3.27 | 0.079 |
| Omoist:Otemp | **7.21** | **0.011** | 0.12 | 0.734 | 1.22 | 0.277 | 0.22 | 0.640 |

Supporting information 3. Pair correlation matrix, based on Pearson ´s correlation coefficient, for dependent variables. Significant values (≤ 0.05) and variables further used for statistical analyses are underlined.

| | Prop. dormant seeds | GI | Total germination | Prop. viable seeds | T50 |
|---|---|---|---|---|---|
| Prop. dormant seeds | 1 | **-0.861** | **-0.987** | **0.419** | **0.737** |
| GI | | 1 | **0.899** | -0.087 | **-0.784** |
| Total germination | | | 1 | -0.267 | **-0.746** |
| Prop. viable seeds | | | | 1 | -0.029 |
| T50 | | | | | 1 |

Supporting information 4. Effect of seed mass, original (O) and target (T) conditions on total germination, proportion of viable seeds and time to 50% germination (T50) assessed using mixed effects models with population used as a random factor. Table A represents results of two boreal and two subboreal populations (the third analysis), Table B represents results of boreal populations (the second analysis). Significant values (≤ 0.05) are in bold. * indicate significant result in the model not including seed mass. ● indicate non-significant results in the model not including seed mass

| **A** | Total germination | | Prop. viable seeds | | T50 | |
|---|---|---|---|---|---|---|
| | F-values | p-value | F-values | p-value | F-values | p-value |
| OMoist | 0.22 | 0.603* | **28.8** | **<0.001** | 0.01 | 0.941* |
| OTemp | 0.72 | 0.400* | 0.01 | 0.919* | 0.70 | 0.400 |
| TMoist | **811.22** | **<0.001** | **14.63** | **<0.001** | 3.36 | 0.070 |
| TTemp | **142.15** | **<0.001** | 0.72 | 0.400 | 0.90 | 0.350 |
| Seed mass | **36.87** | **<0.001** | **15.52** | **<0.001** | 0.05 | 0.928 |
| TMoist:TTemp | **27.59** | **<0.001** | **19.25** | **<0.001** | 0.11 | 0.740 |
| TMoist:OTemp | 0.03 | 0.792 | **8.09** | **0.002** | 1.09 | 0.300 |
| TMoist:Seed mass | 3.06 | 0.058 | 2.67 | 0.075 | 0.78 | 0.380 |
| TMoist:OMoist | 0.48 | 0.502 | **4.67** | **0.024** | 1.18 | 0.280 |
| OMoist:TTemp | **7.47** | **0.006** | 0.14 | 0.208 | 1.75 | 0.190 |
| OMoist:Seed mass | **28.65** | **<0.001** | 0.64 | 0.442 | 2.56 | 0.110 |
| TTemp:OTemp | 0.95 | 0.329 | **5.37** | **0.02** | 0.07 | 0.790 |
| TTemp:Seed mass | **4.60** | **0.015** | 0.27 | 0.598 | 0.14 | 0.710 |
| OTemp:Seed mass | **5.45** | **0.012** | **3.87** | **0.042** | 0.14 | 0.710 |
| TMoist:TTemp:OTemp | 0.01 | 0.929 | 1.96 | 0.158* | 0.43 | 0.520 |
| TMoist:TTemp:Seed mass | 0.11 | 0.814 | **7.20** | **0.003** | 1.40 | 0.240 |
| TMoist:OTemp:Seed mass | 0.49 | 0.511 | 2.32 | 0.126 | 0.64 | 0.420 |
| OMoist:TMoist:TTemp | 0.68 | 0.433* | 2.11 | 0.138 | 0.47 | 0.490 |
| OMoist:TMoist:Seed mass | 0.02 | 0.882 | **6.62** | **0.009** | 2.01 | 0.160 |
| TTemp:OTemp:Seed mass | 0.08 | 0.893 | 2.32 | 0.126 | 0.03 | 0.870 |

| **B** | Total germination | | Prop. viable seeds | | T50 | |
|---|---|---|---|---|---|---|
| | F-values | p-value | F-values | p-value | F-values | p-value |
| TTemp | **43.16** | **<0.001** | 1.90 | 0.320 | 3.07 | 0.083 |
| TMoist | **945.48** | **<0.001** | 0.07 | 0.902 | **13.60** | **<0.001** |
| OMoist | 0.05 | 0.928 | 1.83 | 0.318* | 0.17 | 0.863 |
| Seed mass | 1.57 | 0.672 | 1.54 | 0.398 | 0.19 | 0.847 |
| TTemp: TMoist | **8.21** | **<0.001** | **30.88** | **<0.001** | 2.18 | 0.143 |
| TTemp: OMoist | 2.55 | 0.128 | 0.43 | 0.823 | 3.30 | 0.072 |
| TMoist: OMoist | **10.46** | **<0.001** | **8.26** | **<0.001•** | **6.16** | **0.015•** |
| TTemp: Seed mass | **8.48** | **<0.001** | 2.44 | 0.087 | 2.49 | 0.117 |
| TMoist: Seed mass | **32.31** | **<0.001** | **5.79** | **0.001** | **8.72** | **0.004** |
| OMoist: Seed mass | **14.80** | **<0.001** | **5.89** | **0.018** | 0.02 | 1.000 |
| TTemp: TMoist:OMoist | 2.36 | 0.201 | 1.29 | 0.263 | 2.51 | 0.116 |
| TTemp: OMoist:Seed mass | 1.57 | 0.683 | **9.14** | **0.002** | 0.03 | 0.874 |
| TTemp: TMoist:Seed mass | 0.00 | 0.949 | 0.49 | 0.431 | **7.48** | **0.007** |
| TMoist:OMoist:Seed mass | 0.01 | 0.926 | 0.00 | 0.892 | 0.04 | 0.832 |
